# Guided tokenization and domain knowledge enhance genomic language models’ performance

**DOI:** 10.64898/2026.02.16.706213

**Authors:** Vedant Mahangade, Matthew Mollerus, Keith A. Crandall, Ali Rahnavard

## Abstract

Adapting language models to genomic and metagenomic sequences presents unique challenges, particularly in tokenization and task-specific generalization. Standard methods, such as fixed-length k-mers or byte pair encoding, often fail to preserve biologically meaningful patterns essential for downstream tasks. We introduce *Guided Tokenization* (GT), a strategy that prioritizes biologically and statistically important subsequences based on importance scores, model attention, and class distributions. Combined with *domain adaptation*, which incorporates prior domain specific biological knowledge, this approach improves both representation quality and classification accuracy in compact genomic language models (gLMs). GT enhances biological awareness in genomic language models, particularly for effective small and mid-sized models across key tasks, including DNA sequence read classification, promoter detection, antimicrobial resistance classification, and targeted amplicon taxonomic profiling. Our results highlight the promise of guided tokenization and domain-aware modeling for building efficient, biologically grounded language models for scalable genomic applications.

## 1 Main

In the context of genomic language models (gLMs), the “text” corresponds to biological sequences, such as DNA, RNA, or Amino Acids. Mirroring the paradigm of pre-trained large language models (LLMs), gLMs are trained on a large corpus of genomic data, which must first be tokenized (broken down into small component ‘words’ or tokens). Fine-tuning language models is the standard method for adapting these models to a domain-specific task, involving the partial or full update of the foundational model weights. However, this process does not update the tokenizer, which retains the same vocabulary and merge orders from pre-training. gLMs typically employ tokenization strategies adapted from natural language models, such as k-mer or Byte Pair Encoding (BPE) [1]. However, these algorithms can result in the fragmentation of biologically significant sub-sequences that carry vital information for genomics applications.

For instance, in promoter prediction tasks, one of the most important subsequences is the TATA box, characterized by repeating T and A of 5 to 6 base pairs. This promoter subsequence plays a vital role in regulating transcription by indicating to other molecules where transcription begins [2]. Standard tokenization techniques for a foundational or a fine-tuned model may break such motifs into smaller, biologically irrelevant subtokens (Figure S4a and S4b), ultimately impairing model performance on tasks that depend on the recognition of complete biological patterns. Studies [3] have shown that augmenting and preserving such subsequences to an existing model vocabulary improves the performance of LLMs across various domain-specific tasks. To address this, we introduce Guided Tokenization (GT), a domain-aware tokenization approach that aims to enhance the performance of gLMs on domain-specific tasks. Unlike BPE, which may disrupt biologically significant motifs, GT leverages domain-specific knowledge to preserve key patterns as individual tokens and prioritizes token ID generation for these motifs (Figure S4c). This targeted approach not only improves the accuracy of tasks such as promoter detection but also enables the model to better capture sequence patterns relevant to specific tasks (Figure S2) during inference and fine-tuning.

We developed multiple GT strategies (Figure 1a) to enhance gLM performance on downstream tasks by incorporating biologically meaningful sequence information (Figure 1c). We implement these strategies on pre-trained foundational gLMs, such as *DNABERT2* 117M [4] and *seqLens* 87M [5]. The strategies include: **(1) weighted tokens**, in-vocabulary tokens prioritized based on attribution scores before finetuning; **(2) unique class-specific k-mers**, prioritized based on frequency and length, capturing important subsequences. We evaluated GT against standard BPE fine-tuned models to assess their impact on model accuracy, representation quality, and computational efficiency in genomic sequence analysis by fine-tuning base models across tasks such as promoter detection, antibiotic resistance gene (ARG) classification, and 16S rRNA-based genus identification, each presenting unique biological and computational challenges. We also compare the performance of these models against widely used alignment-based tools such as ResFinder [6] for ARG classification and DADA2 [7] for 16S genera classification.

**Fig. 1:**
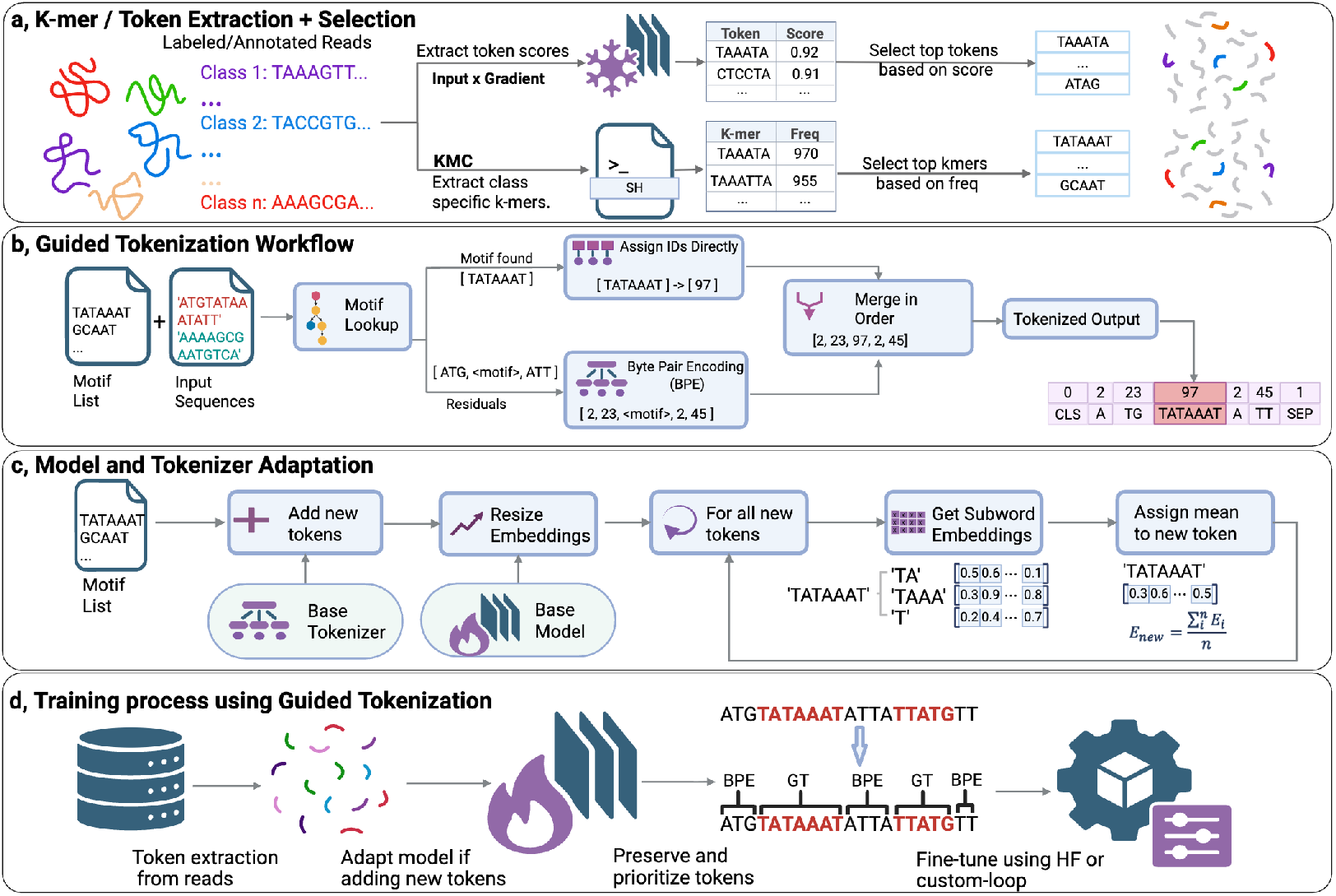
Guided Tokenization for token prioritization augmentation, prioritization, and preservation. **a**, K-mer and token extraction workflow. **b**, Guided tokenization (GT) pipeline with BPE fallback. **c**, Model adaptation procedure using mean subword embeddings. **d**, Training a model using Guided Tokenization.

In promoter vs non-promoter binary classification, models trained with GT (gLM - GT) demonstrated consistent improvements over models trained using BPE tokenization (gLM-BPE). The unique k-mer GT strategy achieved the highest F1 Score (82.88% vs. 78.93% for BPE), with notable gains in recall (81.2% vs. 74.16%) and accuracy (83.69% vs. 80.79%) (Figure 2a). Among test sequences utilizing GT-specific tokens (n=52/613), misclassification rates decreased from 28.85% to 23.08% compared to BPE tokenization of the same sequences (Figure 2b). Probability distribution analysis revealed that GT-tokenized sequences showed increased confidence in correct predictions, with true positives and true negatives concentrated at higher probability thresholds (Figure 2c). Analysis of near-miss predictions showed that GT substantially reduced false negatives in promoter-to-non-promoter transitions, with an average probability margin of 0.3 (Figure 2d).

**Fig. 2:**
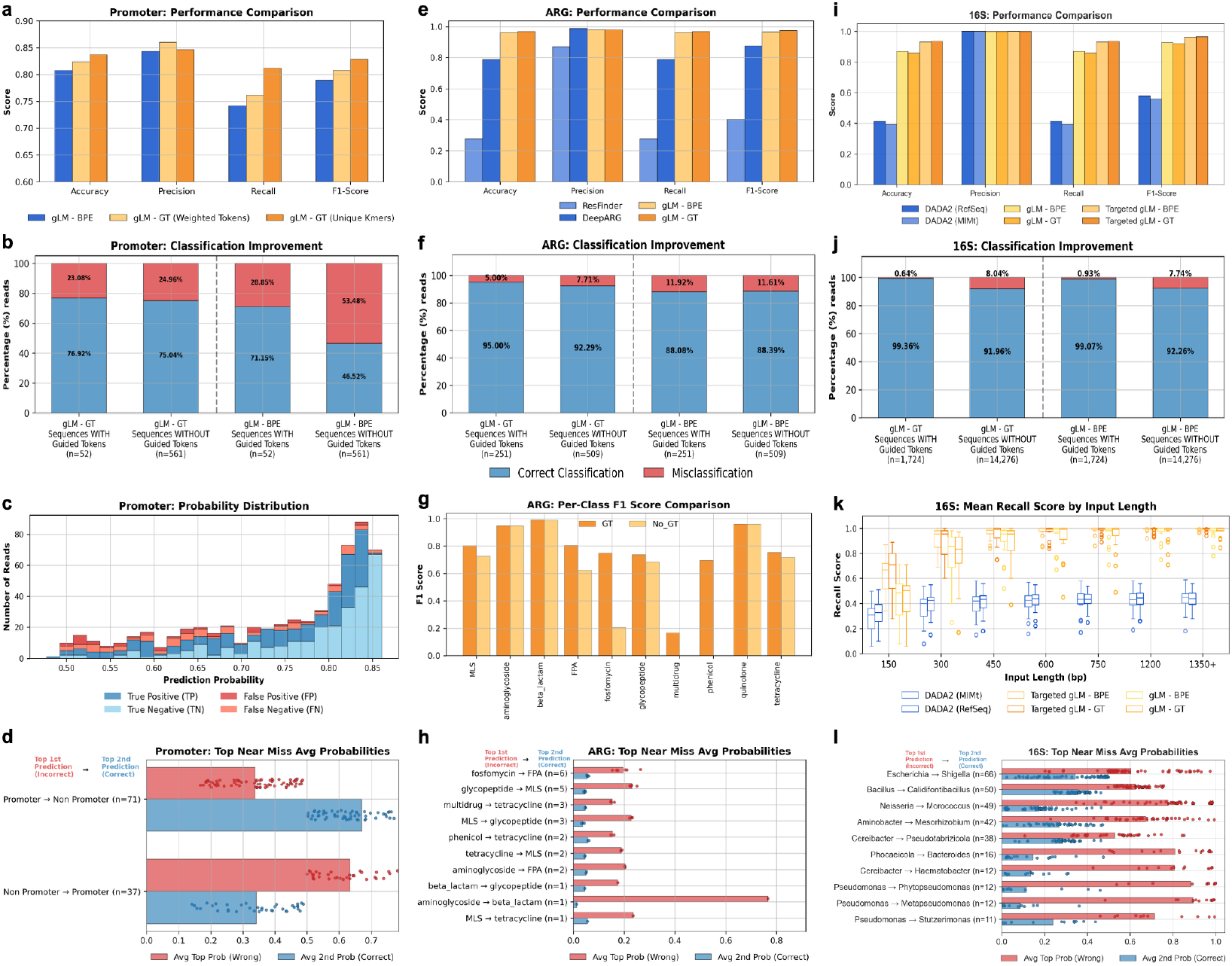
Performance comparison of GT across genomic classification tasks. **a–d**, Promoter sequence classification. **e–h**, Antibiotic Resistant Gene (ARG) classification. **i–l**, 16S rRNA taxonomic classification.

GT achieved marked improvements in multi-class ARG classification (94.48% accuracy vs. 92.28% for BPE), substantially outperforming traditional tools like DeepARG (71.9%) and ResFinder (13.3%) (Figure 2e). Sequences containing GT tokens (n=251/760) demonstrated a 58% reduction in misclassification rate (5.0% vs. 11.92% for BPE) (Figure 2f). Following the trends from promoter vs non-promoter classification, GT improved performance even for sequences not directly utilizing guided tokens, reducing their misclassification rate from 11.61% to 7.71%, suggesting that incorporation of resistance-associated k-mers into the vocabulary enhanced overall model representations. Per-class F1-scores revealed GT’s particular strength in classes with limited training examples (115 training samples for multidrug vs 4211 for beta lactam), where domain-specific k-mers compensated for data scarcity (Figure 2g). Near-miss analysis showed that GT reduced ambiguity between mechanistically related resistance classes, with most confusion limited to beta-lactam subclasses where probability margins remained high (*>* 0.6 bp) (Figure 2h). For binary (ARG vs non-ARG) classification, when compared against BPE, GT not only improves classification accuracy but also yields better-calibrated probability estimates, achieving a lower Brier score of 0.216 vs. 0.224 of BPE (Figure S6b).

The 16S classification task, involving 4,288 genera across 335 orders, revealed GT’s scalability limitations in extremely high-dimensional classification spaces. While gLMs substantially outperformed alignment-based DADA2’s taxonomic assignment (87.1% and 93.0% for GT and BPE vs. 41.3% accuracy), GT showed marginal underperformance relative to BPE in the standard sequence to genus model configuration (85.8% vs. 87.1%) (Figure 2i). However, a hierarchical ensemble approach (Targeted gLM) that first classified sequences at the order level before genus-level classification improved GT performance to 93.47% accuracy, slightly higher than BPE, which achieved 93.06% accuracy(Figure 2j). In this hierarchical framework, GT demonstrated advantages across different sequence lengths, achieving slightly higher mean recall scores (Figure 2k). Sequences utilizing GT tokens (n=1,724/16,000) maintained lower misclassification rates (0.64% vs. 0.93% for BPE), though unlike the previous two tasks, the misclassification rate for non-GT tokenized sequences increased from 7.74% to 8.04%. Evaluation conducted on varying sequence length revealed that the model’s top predictions are more confident (concentrated between 0.9 and 1.0) with longer sequences (*>* 1200 bp). The primary source of error for both approaches was the Escherichia-Shigella distinction (Figure 2l), a known challenge in 16S-based taxonomy due to their close phylogenetic relationship and limited sequence divergence in this marker gene (Figure S10b and c).

Performance analysis across tasks, the primary performance benefits of GT over BPE, were realized by augmenting domain-specific k-mers to the vocabulary and model before fine-tuning. This effectiveness is modulated by the ratio of biological classes to vocabulary capacity. For promoter detection (2 classes) and ARG classification (12 classes), adding 100-500 class-specific k-mers per class (10% - 30% of original vocabulary size) yielded consistent improvements. In contrast, the 16S task’s 4,288-genera constrained per-class token allocation, limiting GT’s ability to capture genus-specific signatures without vocabulary explosion. This suggests that GT is most effective for tasks where biologically meaningful subsequences can be adequately represented within existing vocabulary constraints, or when hierarchical modeling can reduce the effective number of classes per model.

## 2 Online Content

Any methods, additional references, Nature Portfolio reporting summaries, source data, extended data, supplementary information, acknowledgments, peer review information, details of author contributions and competing interests; and statements of data and code availability are available at https://github.com/omicsEye/guidedtokenizer.

## 3 Methods

Guided Tokenization is implemented as a standalone Python script, essentially a class that inherits from the FastTokenizer package of the HuggingFace transformer library. The entire process of implementing the guided tokenizer is divided into three phases: (**1**) important token/k-mer *Extraction*, (**2**) tokenizer and model *Augmentation*, and (**3**) *Fine-Tuning* using the augmented tokenizer and model. Training and evaluation of our GT approach was evaluated on three biologically targeted datasets: (**1**) promoter detection, (**2**) antibiotic drug resistance variants, and (**3**) 16S rRNA gene classification of bacterial diversity. We first describe our three target datasets and then proceed to the algorithm descriptions and the training, validation, and testing using these data.

### 3.1 Datasets

We performed a robust evaluation of tokenization strategies under both controlled and naturally noisy scenarios, which is critical for developing generalizable models in metagenomic and microbial ecology applications.

#### PROMOTER detection

For this task, we used the promoter dataset from the GUE benchmark suite [4]. The dataset comprises approximately 59,000 DNA sequences, each 300 base pairs in length, labeled as either promoter or non-promoter. We use an 80-10-10 split for training, validation, and testing.

#### ARG Classification

This task focused on classifying antibiotic drug classes from sequences. We used the long read ARG dataset from resLens [8] available from Huggingface at https://huggingface.co/datasets/omicseye/long_read_data, which was originally curated from ResFinder [6] and The NCBI Pathogen Detection Project [9]. The dataset contains 15,212 samples with 50% ARG reads and the remaining 50% tagged as non ARG. We filtered the dataset to only include ARG classes, resulting in 7606 sample reads across 12 drug classes, of which 6086 were used for training and 760 for validation and testing each.

#### 16S Taxonomic Classification

This task focused on genus-level classification of partial 16S rRNA sequences. The dataset used was curated for 16S gLM (https://github.com/omicsEye/16S_gLM), and the curation process involved simulating long read sequences using SimLord [10] from the clean RefSeq 16S database [11][12] to account for class imbalance and limited training data per class, given 4288 classes. The final dataset contained 30,000 simulated reads per class for training and 10,000 reads per class for validation and testing each.

### 3.2 Important Token/k-mer *Extraction*

Important tokens or k-mers that can serve as meaningful sequence motifs in the GT framework can be derived in multiple ways. We implemented two complementary strategies that leverage both the training data and the foundation model’s pretrained knowledge to identify subsequences representative of the domain corpus.

#### Weighted Tokens

We applied an input×gradient attribution method [13] across the training dataset to estimate the contribution of each token to the model’s predic-tions. This saliency analysis highlights which vocabulary tokens the pretrained model relies on when making correct zero-shot predictions. After filtering for correctly predicted samples, we retained only tokens with log-normalized scores above a threshold determined using the elbow method [14][15]. While the elbow criterion is a heuristic and has known limitations in clustering [16], in this setting, it serves solely to identify a cutoff in the ranked attribution scores. We verified that the qualitative set of informative subsequences is stable across a reasonable range of thresholds(Figure S1). This procedure identifies informative subsequences already present in the model’s existing vocabulary (Figure 1a).

#### Unique k-mers

Using KMC [17], we extracted all possible k-mers (k = 5–25) from the annotated training data and then identified class-specific unique k-mers (Figure 1a). The number of unique k-mers per class depends on sequence length and class size. Longer sequences yield more distinct k-mers, whereas larger class sizes reduce the uniqueness count. For the promoter dataset, roughly 10 million unique k-mers were identified per class (Figure S8a). In the multi-class ARG dataset, unique k-mer counts reflected class imbalance, with beta lactam generating the largest set (∼7 million; Figure S9a). From these unique sets, we selected the highest-frequency k-mers for vocabulary augmentation: the top 500 per class for the promoter task and the top 100 per class for ARG. These thresholds were chosen to balance biological coverage and vocabulary efficiency, corresponding to approximately 10–30% vocabulary expansion, and also aligning with observations that overly large vocabularies can harm generalization and downstream performance in genomic language models such as seqLens [5]. By prioritizing recurrent, class-informative motifs and avoiding rare, longtail k-mers, we maintain a compact vocabulary that supports both performance and computational tractability. For the 16S dataset, where the number of genera (4,288) exceeds the model’s vocabulary capacity (4,096), we instead selected the top k-mers at the taxonomic order level, producing 1,790 high-frequency k-mers. This approach yields only 3–5 k-mers per order; to address this limitation, we adopted a hierarchical ensemble architecture, reducing the number of classes processed per model and enabling the incorporation of additional k-mers. All k-mer extraction steps were performed exclusively on the reference reads used for generating training data to prevent data leakage. Finally, all selected tokens and k-mers were sorted by decreasing subsequence length following the Long Token First principle [18], which suggests that longer subsequences tend to encode richer semantic or functional information.

### 3.3 Model and Tokenizer *Augmentation*

The weighted tokens are selected using gradient-based attribution. These tokens already exist in the tokenizer vocabulary and therefore do not require changes to the vocabulary and their embeddings. In contrast, the unique k-mer approach identifies out-of-vocabulary (OOV) subsequences that must be added to the tokenizer’s vocabulary to fully capture the domain-specific sequence patterns. Incorporating these new k-mers requires expanding the model’s embedding layer. By default, this expansion initializes the embeddings for newly added tokens randomly. Although fine-tuning eventually updates these parameters, starting from random initialization prevents the model from leveraging the pretrained representations learned from the foundational corpus. To avoid this limitation, we initialize the embeddings of newly added k-mers using the mean of their constituent subword embeddings, following the efficient domain-adaptation strategy described by Sachidananda et al. [19]. This initialization anchors new tokens within the pretrained embedding space, enabling more effective transfer of the model’s prior knowledge.

#### Algorithm 1

Model and Tokenizer Augmentation with Mean Subword Initialization

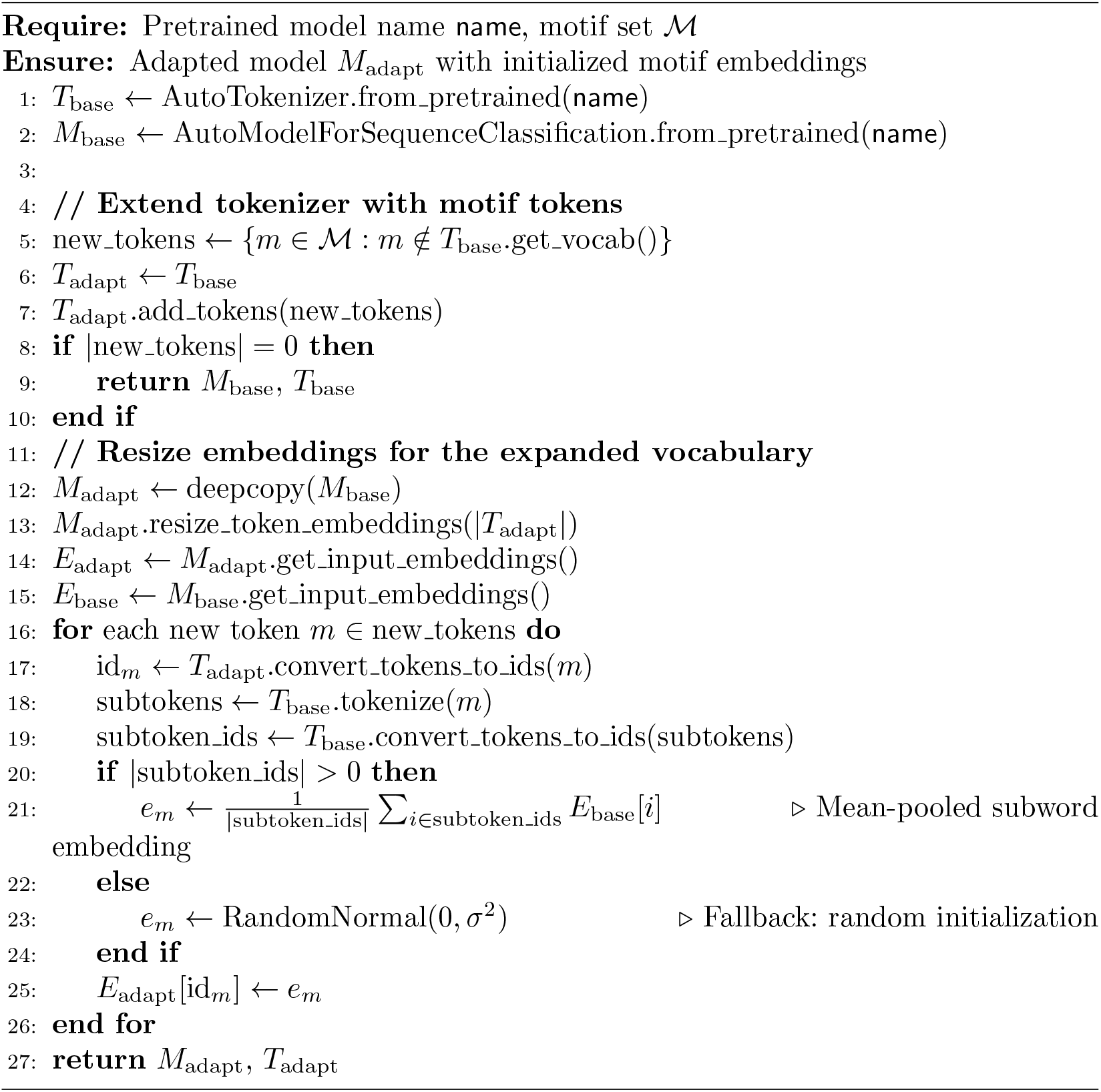

### 3.4 Guided Tokenization

We implemented a GT approach to preserve important motifs during the tokenization process. The function acts as a wrapper around the pre-built BPE tokenizer. The algorithm operates in three phases: **(1)** construction of a trie data structure [20] from a predefined motif set for efficient pattern matching, **(2)** linear-time motif detection across the input sequence using the trie datastructure, and **(3)** hybrid tokenization where detected motifs are preserved as single tokens while intervening sequences are processed by the base BPE tokenizer. The algorithm achieves O(n) time complexity for motif detection, where n is the sequence length, by traversing the input once while simultaneously matching against all motifs via the trie structure (Figure S3). Nonoverlapping motifs are prioritized, with the algorithm advancing past each detected motif to prevent redundant matches. This approach maintains the semantic integrity of functional elements that conventional subword tokenization might fragment. While trie structure provides efficient motif lookup, the process adds a minimal overhead in tokenization time when compared to BPE (Figure S5b). However, GT is substantially faster than k-mer tokenization, which produces more tokens due to the overlap of k-1. Two operational modes are supported: **augment mode**, which expands the vocabulary with novel motif tokens, and **prioritize mode**, which leverages existing motif tokens in pre-trained vocabularies without modification. This flexibility enables seamless integration with existing genomic language models while preserving domain-specific biological knowledge.

#### Algorithm 2

Guided Tokenization with Motif Preservation

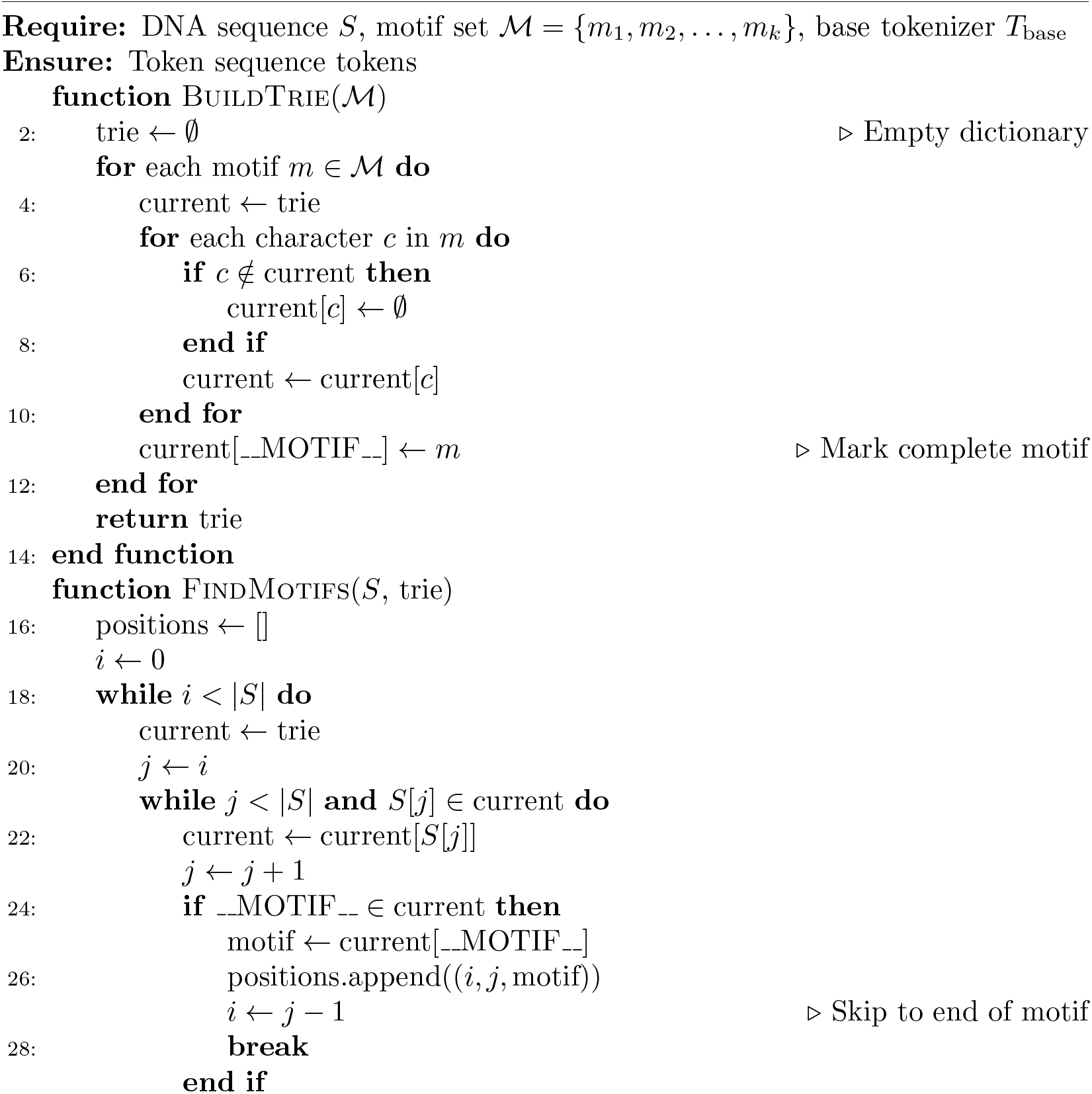
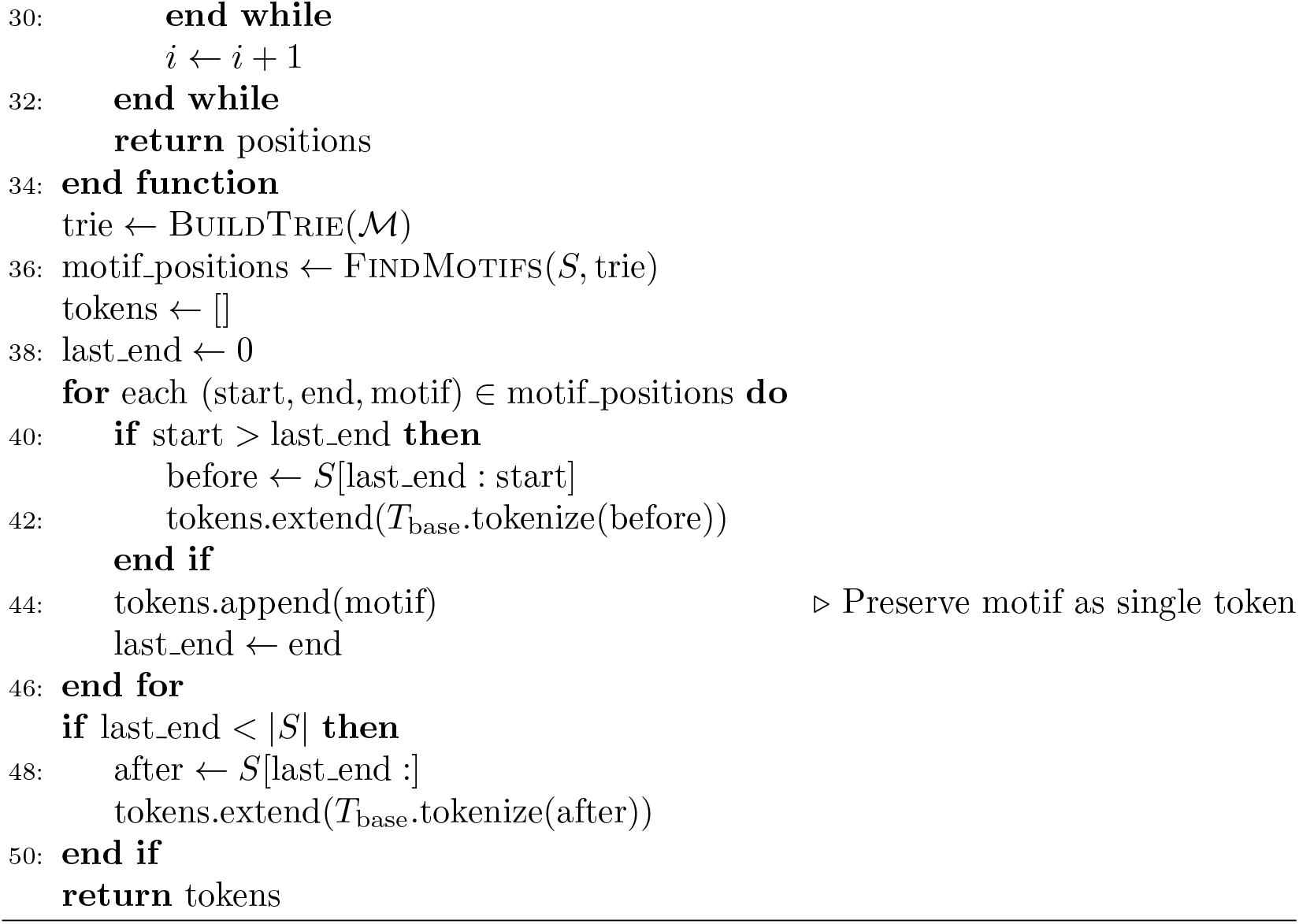

### 3.5 *Fine-Tuning* and Evaluation

We used BPE and GT to fine-tune base models for each task in order to conduct a thorough evaluation. Every experiment was performed using SLURM on GPU nodes with 1-4 NVIDIA Tesla V100 SXM2 16GB GPUs on the George Washington University High Performance Computing cluster. Single GPU nodes were used to train models for promoter and ARG tasks. Due to the large number of training samples for 16S classification (∼128 million), we used distributed computing via data parallelization across 4 GPUs.

#### Promoter vs Non-Promoter Classification

We first fine-tuned DNABERT2 on a binary promoter–non-promoter classification task to compare two GT strategies against the standard BPE tokenizer, which serves as our gLM baseline. For the weighted-token strategy, we prioritized 498 subsequences with log-normalized importance scores above the elbow-method threshold (–2.65; Figure S1). For the unique k-mer strategy, we selected the top 500 k-mers from promoter sequences and the top 500 from non-promoter sequences. These subsequences were added to both the model vocabulary and tokenizer. In both approaches, subsequences were sorted in decreasing order of length to ensure longer k-mers received higher priority. DNABERT2 was fine-tuned for 10 epochs using a learning rate of 3e-5, batch size of 8, and weight decay of 0.1—matching the hyperparameters reported in Zhou et al. [4]. Evaluation on a held-out test set of 5,900 sequences (Figure 2a; Table S1) showed that the GT model using unique k-mers (gLM–GT unique) achieved the best overall performance. All subsequent experiments, therefore, adopt the GT unique k-mer approach.

#### ARG Classification

To examine the impact of vocabulary augmentation on a different architecture and dataset, we adapted the GT strategy for antibiotic resistance gene (ARG) classification using the seqLens-based 89M model [5], following the methodology of resLens. We compared BPE-based and GT gLMs with two established tools: ResFinder (alignment-based) and DeepARG (alignment plus deep learning) [21]. For the GT model, we extracted all 5–25-mer subsequences from training reads and identified k-mers unique to each class. The top 100 high-frequency k-mers per class (1,200 total) were incorporated into the tokenizer and prioritized by length. Both BPE and GT models were fine-tuned for 10 epochs using a learning rate of 1e-5 and weight decay of 0.1. An effective batch size of 16 was obtained using a batch size of 8 and gradient accumulation steps of 2. ResFinder and DeepARG were run using default settings. Unlike gLMs, which output class probabilities for all categories, ResFinder and DeepARG use alignment to first filter out non-ARG reads; any predictions outside the predefined class set were excluded from evaluation. As shown in Figure 2e and Table S2, the gLM - GT consistently outperformed both BPE and the alignment-based tools across all metrics except precision, for which DeepARG achieved the highest score.

#### 16S Taxonomic Classification

This evaluation focuses on real-world scenarios where the number of classes is exceptionally high. With the number of classes exceeding the existing vocabulary size, implementing GT by augmenting unique k-mers is a challenge. For this task, our dataset consisted of 4288 genera. As mentioned earlier, to implement GT without exceeding 10-30% vocabulary augmentation limit, we selected top k-mers per order instead of genera. This approach only results in 3 to 5 k-mers per order to be added to the model and tokenizer. To overcome this, we limited the number of classes a model can process by creating a hierarchical classifier that leverages lineage from order level to genus level. A single order-level model first assigns each sequence to one of 335 orders. Based on this prediction, the sequence is passed to one of 12 order-specific genus classifiers, each trained on genera sharing a common taxonomic lineage. This hierarchical design constrains the prediction space and improves interpretability by aligning the model architecture with established microbial taxonomy. For each model, we implement GT by augmenting the model and tokenizer with the top 1300 unique k-mers corresponding to their specific class set. Since the taxonomic lineage is not balanced, with more genera in certain orders than others, the number of classes per model was also different. These hierarchical classifiers are referred to as Targeted gLM - GT and Targeted gLM - BPE for GT and BPE variants, respectively, while the non-hierarchical sequence to genus classifiers are referred to as gLM - GT and gLM - BPE. Each model was trained on 100,000 steps with a learning rate of 1e-4. We used a batch size of 32 with gradient accumulation steps of 16 across 4 GPUs, resulting in an effective batch size of 1024. For evaluation, we randomly sampled reads from the held-out dataset with varying lengths (150 bp, 300 bp, 450 bp, 600 bp, 750 bp, 1200 bp, 1350+ bp). We also compared the performance against alignment-based DADA2 [7] with two different reference databases, RefSeq [11] and MiMt [22], which were cleaned by removing nested sequences, duplicates, and naming ambiguities [12]. While gLMs outperform alignment-based DADA2, gLM-GT has lower scores than its BPE counterpart. Figure2i depicts that most performance gain is achieved using hierarchical classification in Targeted gLMs; however, even with limiting the prediction space per model, the Targeted gLM - BPE still showed marginally better scores across metrics compared to Targeted gLM - GT Table S3.

## 4 Data availability

Data availability The models and datasets used in this study are subject to intellectual property protection and are therefore not publicly available. Access may be granted by the corresponding author (A.R.) upon request and subject to appropriate licensing or material transfer agreements.

## 5 Code availability

All of the code used to implement Guided Tokenization, train the models, perform inference, and benchmark may be granted by the corresponding author (A.R.) upon request and subject to appropriate licensing or material transfer agreements.

## 6 Acknowledgments

This work was supported by the National Science Foundation under Grants No. 2109688 and 2507498 awarded to A.R. and K.A.C., and by a Technology Maturation Award from the Technology Commercialization Office at The George Washington University.

## 7 Author contributions

A.R. conceived and designed the methodology. V.M. implemented the approach and performed performance evaluations. V.M., M.M., and A.R. designed the applications and curated the datasets. A.R. and K.A.C. guided data interpretation and structured the results. V.M. and A.R. drafted the manuscript. A.R. and K.A.C. acquired funding for the project. All authors reviewed, discussed, and approved the final manuscript.

## 9 Additional information

**Correspondence and requests for materials** should be addressed to Ali Rahnavard.

## Appendix A

Supplementary Materials

**Table S1:** Test scores for promoter vs non-promoter task across models.

1 Appendix A Supplementary Materials
| Model | Loss | Accuracy | F1 | MCC | Precision | Recall |
| --- | --- | --- | --- | --- | --- | --- |
| gLM – BPE | 0.4169 | 0.8075 | 0.7893 | 0.6177 | 0.8435 | 0.7416 |
| gLM – GT (Weighted Tokens) | 0.4032 | 0.8238 | 0.8078 | 0.6503 | **0.8598** | 0.7617 |
| gLM – GT (Unique K-mers) | **0.3974** | **0.8369** | **0.8288** | **0.6736** | 0.8462 | **0.8121** |

**Table S2:** Test scores for ARG Classification Task.

| Model | Accuracy | Precision | Recall | F1-Score |
| --- | --- | --- | --- | --- |
| ResFinder | 0.0710 | 0.7008 | 0.0710 | 0.1263 |
| DeepARG | 0.6481 | **0.9516** | 0.6481 | 0.7649 |
| gLM - BPE | 0.6979 | 0.8692 | 0.6979 | 0.7527 |
| gLM - GT | **0.7215** | 0.9341 | **0.7215** | **0.7848** |

**Table S3:** Test scores for 16S Classification task.

| Model | Accuracy | Precision | Recall | F1-Score |
| --- | --- | --- | --- | --- |
| DADA2 (RefSeq) | 0.4130 | **1.0000** | 0.4130 | 0.5793 |
| DADA2 (MiMt) | 0.3937 | 0.9998 | 0.3937 | 0.5571 |
| gLM – BPE | 0.8705 | 0.9987 | 0.8705 | 0.9270 |
| gLM – GT | 0.8580 | 0.9993 | 0.8580 | 0.9185 |
| Targeted gLM – BPE | 0.9299 | 0.9998 | 0.9299 | 0.9630 |
| Targeted gLM – GT | **0.9341** | 0.9995 | **0.9341** | **0.9651** |

**Table S4:** Comparison of tokenization strategies and their properties.

| Tokenization | Adaptive | Sequence Compression | Interpretability | Example Models |
| --- | --- | --- | --- | --- |
| Nucleotide Level | No | No | Yes | HyenaDNA, EVO2 |
| K-mer | No | No | Yes | Nucleotide Transformer, DNABERT |
| BPE | Yes | Yes | Yes | seqLens, DNABERT2 |
| GT | Yes | Yes | Yes | Any BPE-based model |

**Fig. S1:**
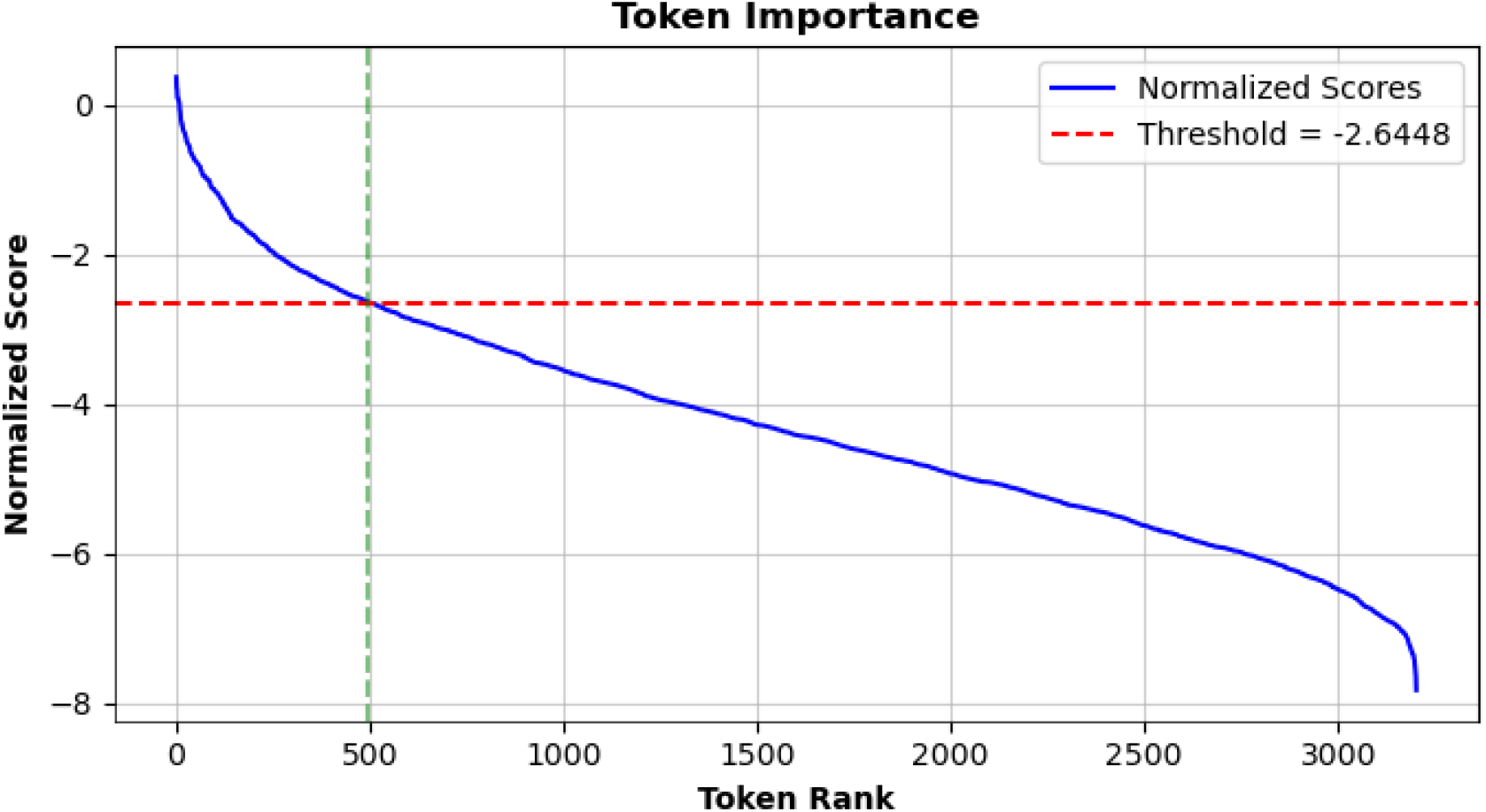
Attribution score by token rank on promoter dataset. Scores are obtained using DNABERT2 on promoter dataset. The elbow method is applied to log-normalized scores to obtain a threshold for token filtering.

**Fig. S2:**
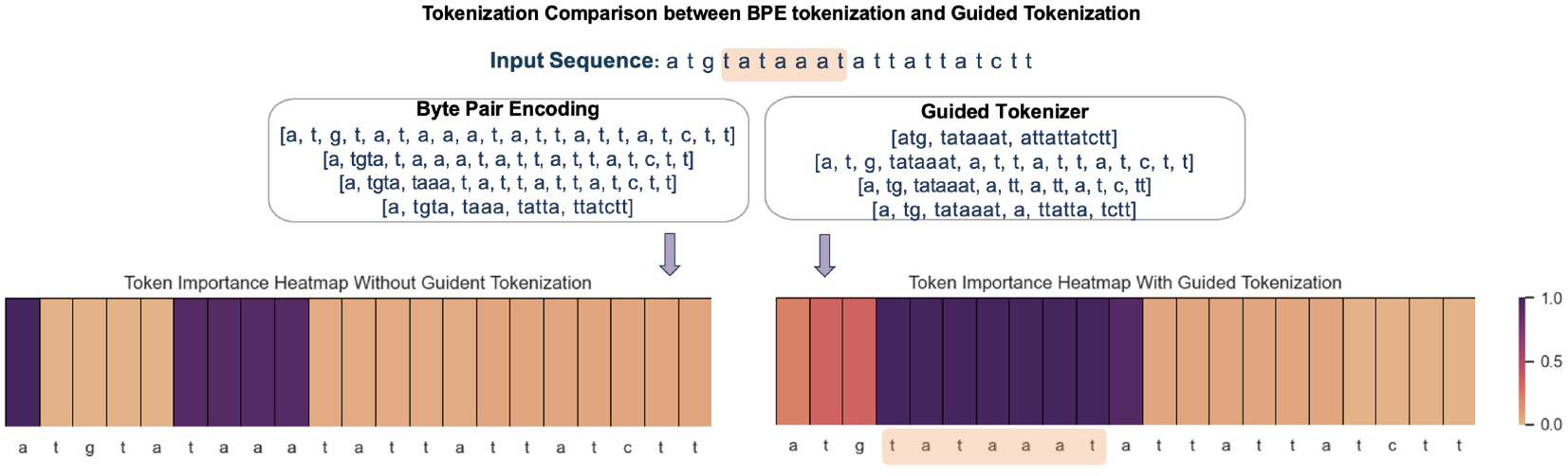
Comparison of BPE and GT: Comparison of tokenization between BPE and GT for the same input sequence. GT preserves meaningful subsequences (e.g., motifs), resulting in a more localized and interpretable heatmap of token importance.

**Fig. S3:**
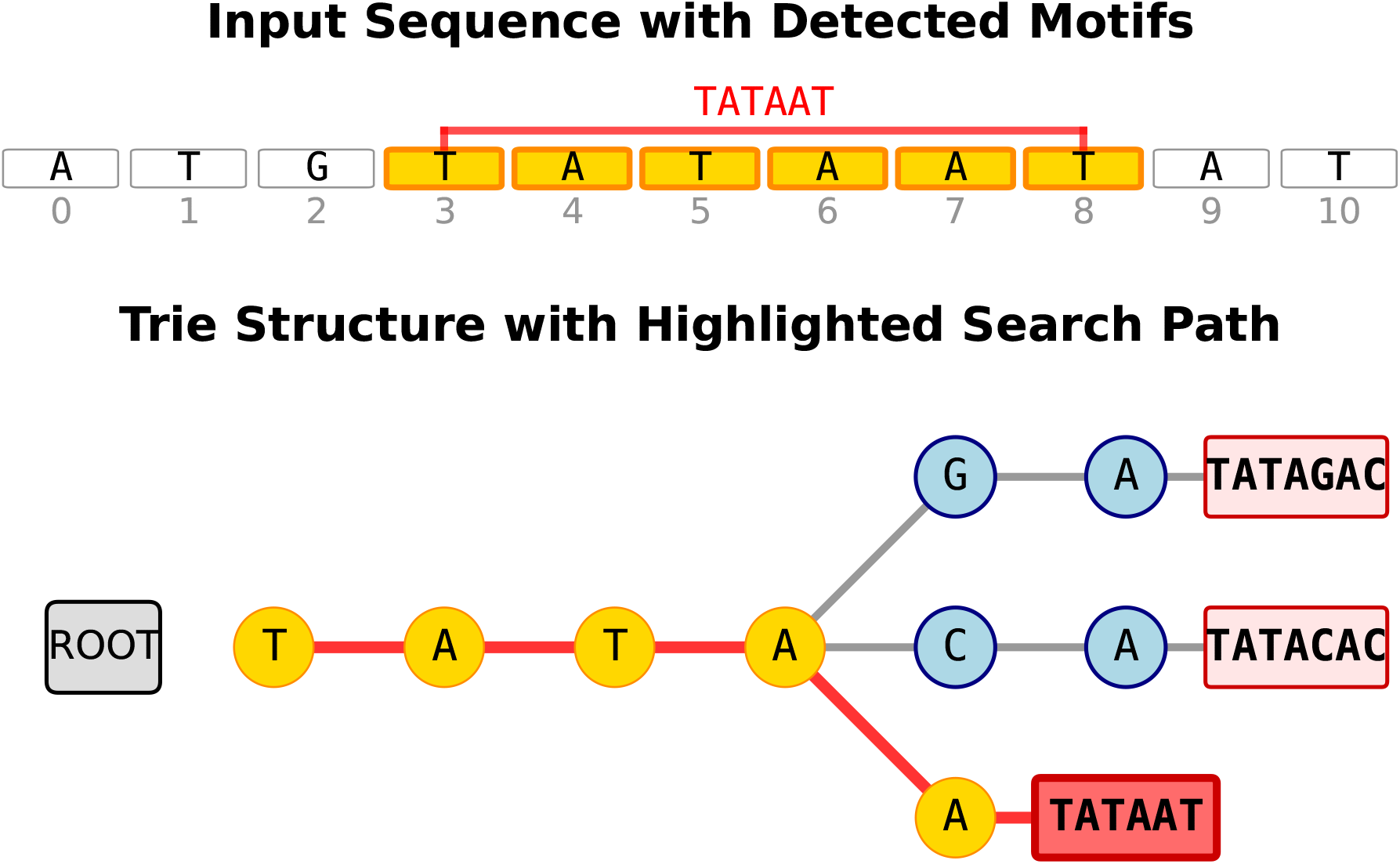
Trie-based motif search: the input sequence with detected TATAAT motif, the trie structure highlights the traversal path used to identify matching patterns.

**Fig. S4:**
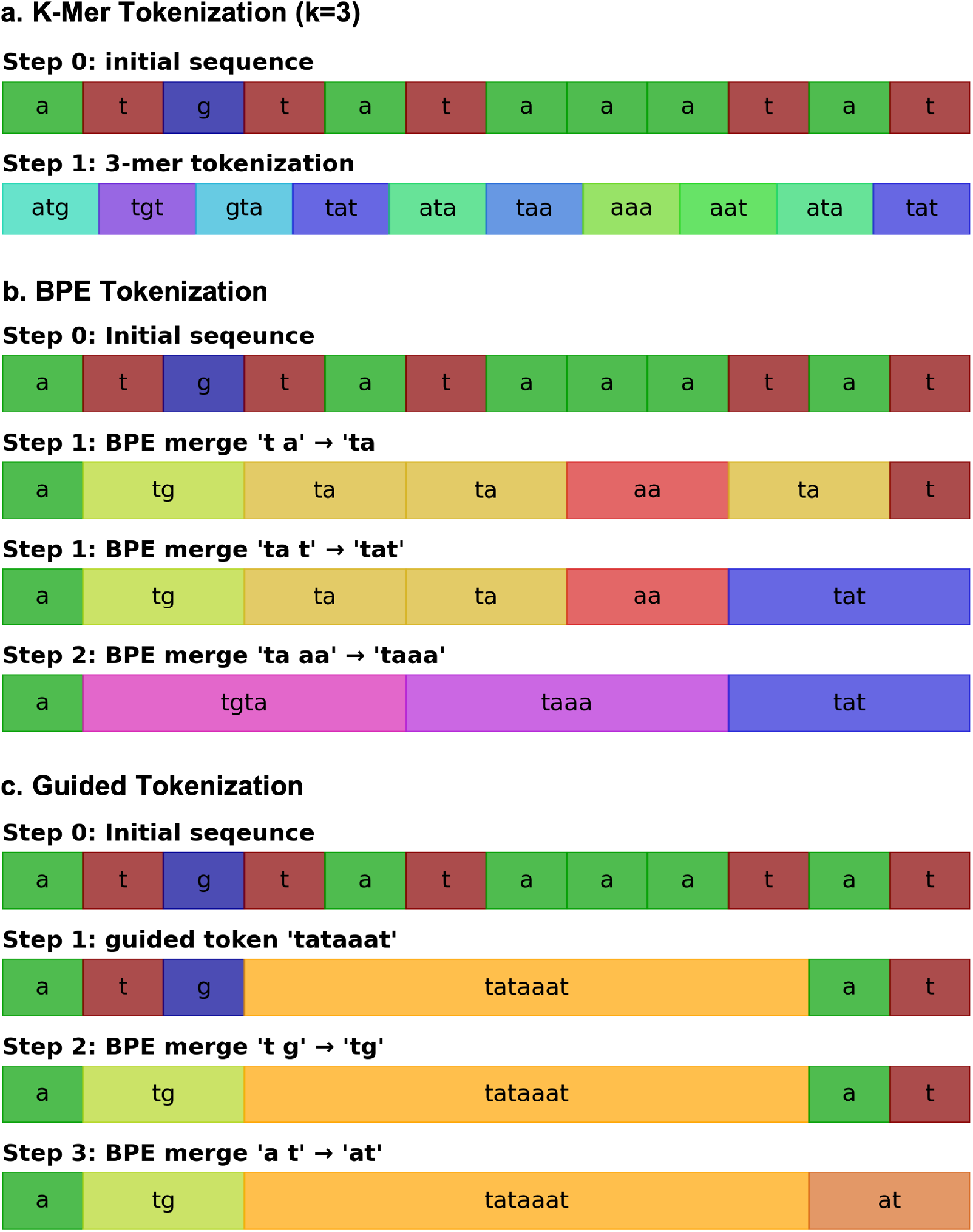
Tokenization schemes: **a. K-mer tokenization** with a k value of 3 generates tokens of uniform length with an overlap of k-1. **b. Byte Pair Encoding (BPE)** demonstrates the sequential process of BPE, showcasing initial sequences and subsequent merges. **c. Guided tokenization (GT)** outlines a structured approach involving prioritization of certain tokens, which preserves them from being broken down by BPE.

**Fig. S5:**
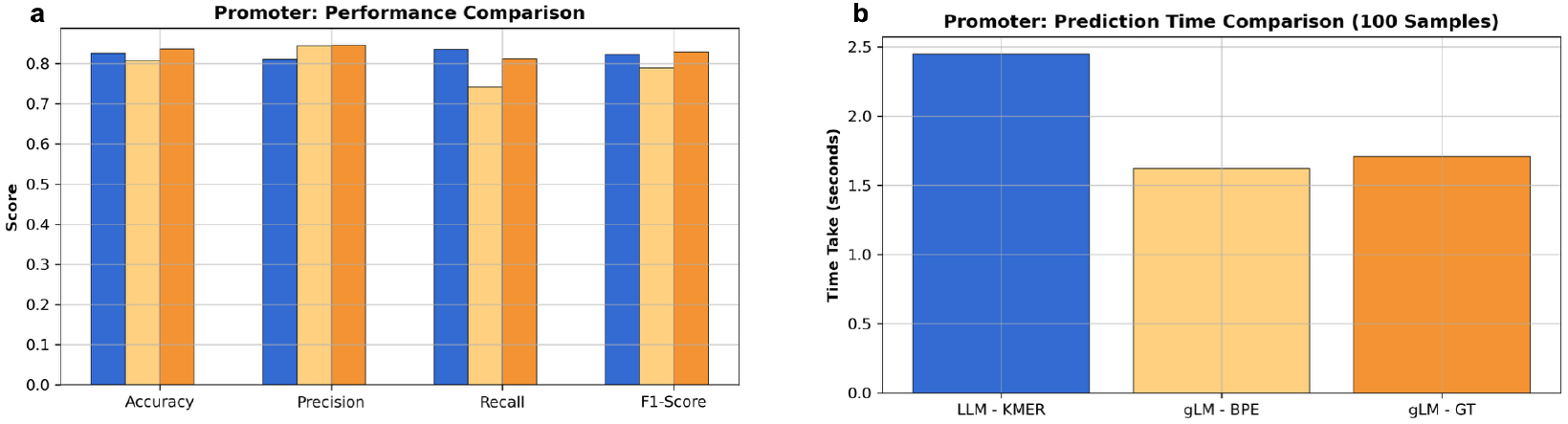
Promoter classification performance and inference efficiency by tokenization methods. **a**, Accuracy, precision, recall, and F1-score for K-MER, BPE, and GT. GT yields the highest F1-score and recall, indicating improved detection of true promoter sequences without sacrificing precision. **b**, Inference-time comparison for promoter prediction across tokenization strategies. GT and BPE achieve inference efficiency comparable to fixed k-mer tokenization. GT adds minimal tokenization overhead during motif preservation when comparedto BPE.

**Fig. S6:**
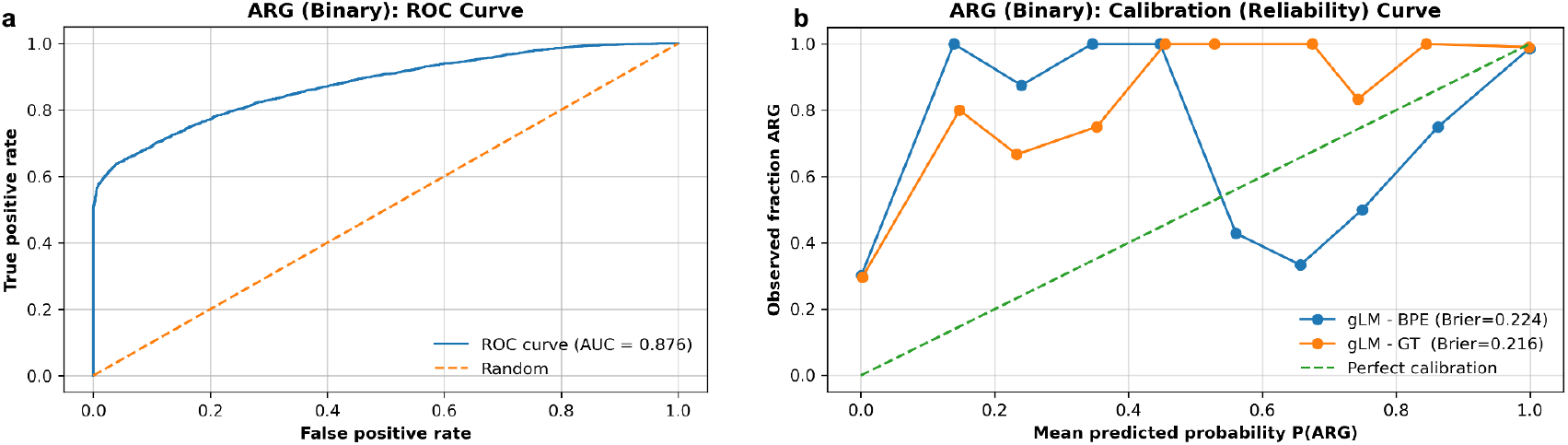
Binary antibiotic resistance gene (ARG) classification performance. **a**, Receiver operating characteristic (ROC) curve for binary ARG detection using gLMs, showing strong discriminative performance with an area under the curve (AUC) of 0.876. **b**, Calibration (reliability) curves comparing gLM–BPE and gLM–GT. GT yields improved probability calibration across confidence bins, reflected by a lower Brier score, indicating more reliable probabilistic outputs for ARG presence prediction.

**Fig. S7:**
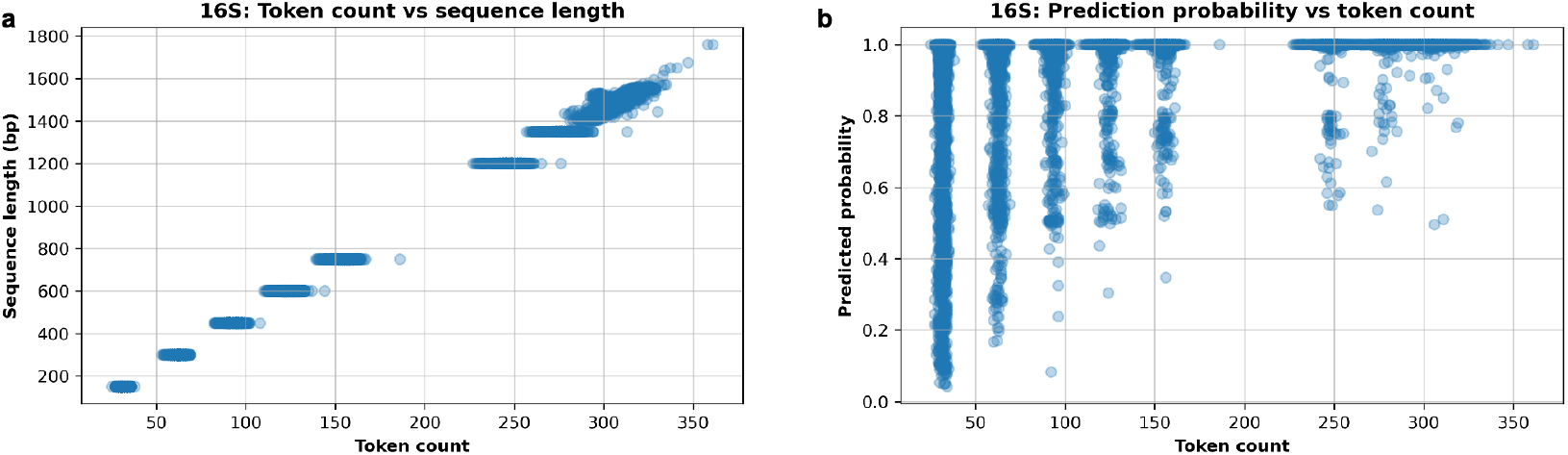
Tokenization characteristics and prediction confidence in 16S classification. **a**, Relationship between input sequence length and resulting token count across simulated 16S reads. Token count increases approximately linearly with sequence length, with discrete clusters corresponding to fixed read-length groups used during evaluation. **b**, Predicted class probability as a function of token count. Longer, more richly tokenized sequences generally yield higher prediction confidence, whereas shorter reads exhibit greater uncertainty, highlighting the impact of effective sequence length on model confidence in high-class-cardinality settings.

**Fig. S8:**
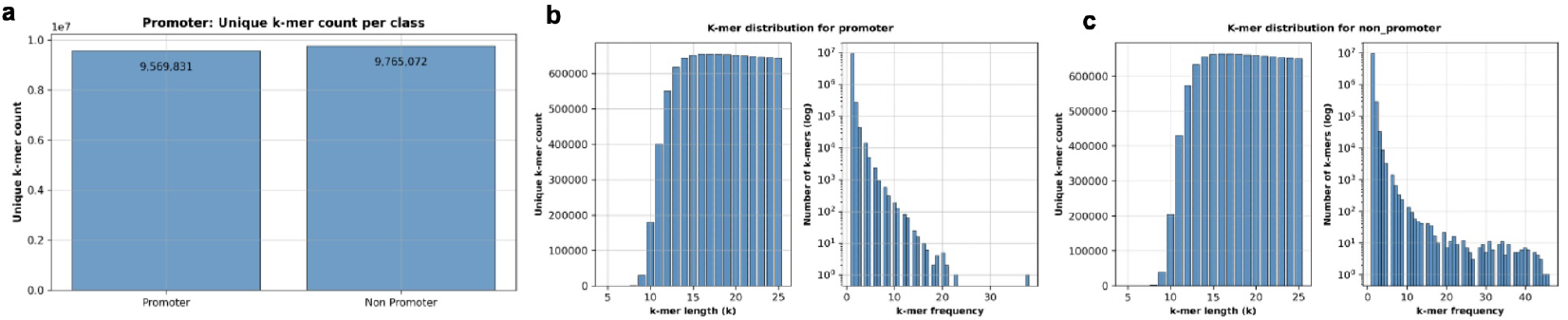
k-mer distribution for promoter vs non-promoter sequences. **a**, Number of unique class-specific k-mers extracted from promoter and non-promoter sequences across k values ranging from 5 to 25, showing comparable k-mer diversity between the two classes despite their distinct functional roles. **b**, k-mer distribution for promoter sequences, showing the number of unique k-mers as a function of k-mer length (left) and the corresponding k-mer frequency histogram on a logarithmic scale (right). Longer k-mers dominate the unique motif space but occur at lower frequencies. **c**, k-mer distribution for non-promoter sequences, exhibiting a similar length–frequency trade-off, with differences in frequency structure reflecting background genomic composition rather than promoter-specific motifs.

**Fig. S9:**
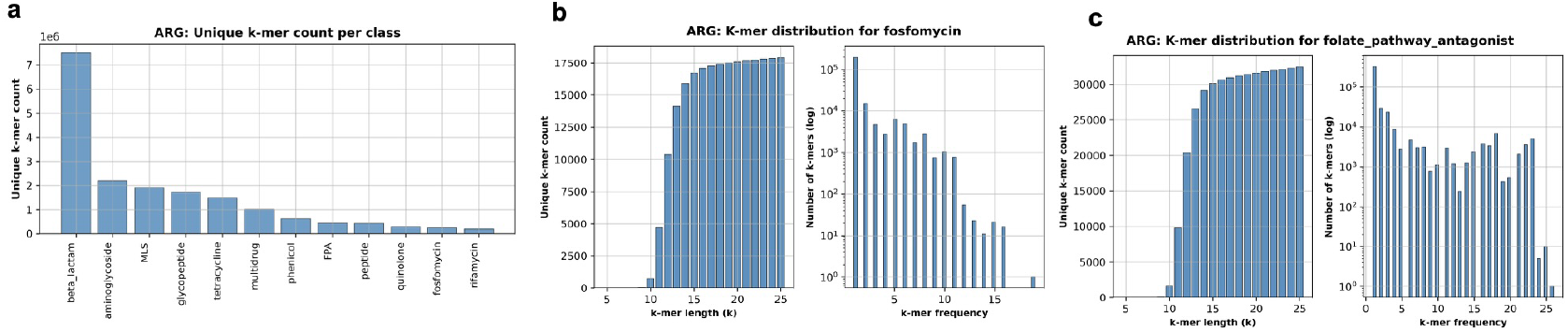
k-mer distribution for ARG classes. **a**, Unique k-mers per antibiotic class obtained for k value 5 to 25 in the training data, with beta-lactam resistance genes yielding the largest unique k-mer set. **b**, k-mer distribution for the fosfomycin resistance class, showing unique k-mer counts by length (left) and the corresponding k-mer frequency histogram on a logarithmic scale (right). **c**, k-mer distribution for the folate pathway antagonist (FPA) resistance class, presented in the same format. Fosfomycin and FPA represent the most frequent near-miss pair in ARG classification, where misclassified sequences most often receive the second-highest probability for the alternate class, highlighting shared or overlapping sequence motifs between these resistance mechanisms.

**Fig. S10:**
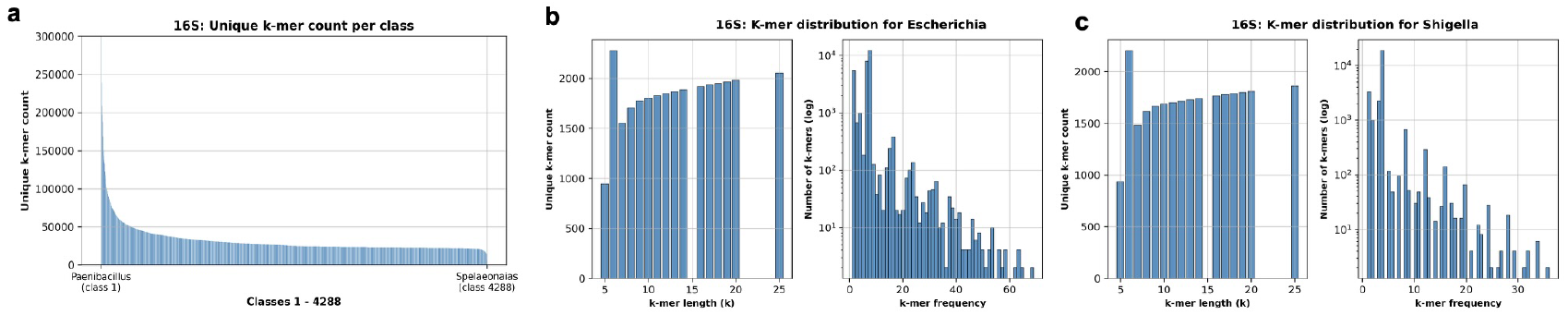
k-mer distribution for 16S sequences by genera. **a**, Number of unique k-mers per genus extracted across k values from 5 to 25 for all 4,288 genera in the dataset, showing a highly skewed distribution in k-mer diversity across taxonomic classes. **b**, k-mer distribution for Escherichia, showing unique k-mer counts by length (left) and k-mer frequency histogram on a logarithmic scale (right). **c**, k-mer distribution for Shigella in the same format. Escherichia and Shigella form the most frequent near-miss pair in 16S classification, reflecting their close phylogenetic relationship and limited sequence divergence within the 16S marker gene.

## References

[1] Schmidt, C. W. et al. Tokenization is more than compression (2024). URL https://arxiv.org/abs/2402.18376.2402.18376.

[2] Oubounyt, M., Louadi, Z., Tayara, H. & Chong, K. T. Deepromoter: Robust promoter predictor using deep learning. Frontiers in Genetics Volume 10 - 2019 (2019). URL https://www.frontiersin.org/journals/genetics/articles/10.3389/fgene.2019.00286.

[3] Zhang, R. et al. Webber, B., Cohn, T., He, Y. & Liu, Y. (eds) Multi-stage pretraining for low-resource domain adaptation. (eds Webber, B., Cohn, T., He, Y. & Liu, Y.) Proceedings of the 2020 Conference on Empirical Methods in Natural Language Processing (EMNLP), 5461–5468 (Association for Computational Linguistics, Online, 2020). URL https://aclanthology.org/2020.emnlp-main.440/.

[4] Zhou, Z. et al. Dnabert-2: Efficient foundation model and benchmark for multispecies genome (2023). 2306.15006.

[5] Baghbanzadeh, M., Mann, B., Crandall, K. A. & Rahnavard, A. seqlens: optimizing language models for genomic predictions. bioRxiv (2025). URL https://www.biorxiv.org/content/early/2025/03/14/2025.03.12.642848.

[6] Florensa, A. F., Kaas, R. S., Clausen, P. T. L. C., Aytan-Aktug, D. & Aarestrup, F. M. Resfinder – an open online resource for identification of antimicrobial resistance genes in next-generation sequencing data and prediction of phenotypes from genotypes. Microbial Genomics 8, 000748 (2022). URL https://app.dimensions.ai/details/publication/pub.1144941138. 10.1099/mgen.0.000748.

[7] Callahan, B. J. et al. DADA2: High-resolution sample inference from illumina amplicon data. Nat Methods 13, 581–583 (2016).

[8] Mollerus, M., Dittmar, K., Crandall, K. A. & Rahnavard, A. reslens: genomic language models to enhance antibiotic resistance gene detection. bioRxiv (2025). URL https://www.biorxiv.org/content/early/2025/07/11/2025.07.08.663767.

[9] National Center for Biotechnology Information. The ncbi pathogen detection project. https://www.ncbi.nlm.nih.gov/pathogens/ (2016).

[10] Stöcker, B. K., Köster, J. & Rahmann, S. Simlord: Simulation of long read data. Bioinformatics 32, 2704–2706 (2016). URL 10.1093/bioinformatics/btw286.

[11] Goldfarb, T. et al. Ncbi refseq: reference sequence standards through 25 years of curation and annotation. Nucleic Acids Research 53, D243–D257 (2025).

[12] Baghbanzadeh, M., Mahangade, V., Crandall, K. A. & Rahnavard, A. Curating 16s rrna databases enhances taxonomic accuracy and computational efficiency in microbial profiling. bioRxiv (2025). URL https://www.biorxiv.org/content/early/2025/11/05/2025.11.04.686545.

[13] Shrikumar, A., Greenside, P. & Kundaje, A. Learning important features through propagating activation differences (2019). URL https://arxiv.org/abs/1704.02685.1704.02685.

[14] Thorndike, R. L. Who belongs in the family? Psychometrika 18, 267–276 (1953). URL https://api.semanticscholar.org/CorpusID:120467216.

[15] Satopaa, V., Albrecht, J., Irwin, D. & Raghavan, B. Finding a “kneedle” in a haystack: Detecting knee points in system behavior, 166–171 (2011).

[16] Schubert, E. Stop using the elbow criterion for k-means and how to choose the number of clusters instead. SIGKDD Explor. Newsl. 25, 36–42 (2023). URL 10.1145/3606274.3606278.

[17] Kokot, M., Dlugosz, M. & Deorowicz, S. Kmc 3: counting and manipulating kmer statistics. Bioinformatics 33, 2759–2761 (2017). URL 10.1093/bioinformatics/btx304.

[18] Lian, H. et al. Lbpe: Long-token-first tokenization to improve large language models (2024). URL https://arxiv.org/abs/2411.05504.2411.05504.

[19] Sachidananda, V., Kessler, J. & Lai, Y.-A. Moosavi, N. S. et al. (eds) Efficient domain adaptation of language models via adaptive tokenization. (eds Moosavi, N. S. et al.) Proceedings of the Second Workshop on Simple and Efficient Natural Language Processing, 155–165 (Association for Computational Linguistics, Virtual, 2021). URL https://aclanthology.org/2021.sustainlp-1.16/.

[20] Bodon, F. & Rónyai, L. Trie: An alternative data structure for data mining algorithms. Mathematical and Computer Modelling 38, 739–751 (2003). URL https://www.sciencedirect.com/science/article/pii/0895717703900586. Hungarian Applied Mathematics.

[21] Arango-Argoty, G. et al. Deeparg: a deep learning approach for predicting antibiotic resistance genes from metagenomic data. Microbiome 6, 23 (2018). URL 10.1186/s40168-018-0401-z.

[22] Cabezas, M. P., Fonseca, N. A. & Muñoz-Mérida, A. Mimt: a curated 16s rrna reference database with less redundancy and higher accuracy at specieslevel identification. Environmental Microbiome 19, 88 (2024). URL 10.1186/s40793-024-00634-w.

